# DAP-seq Reveals Cluster-Situated Regulator Control of Numerous *Streptomyces* Natural Product Biosynthetic Genes

**DOI:** 10.64898/2026.01.05.697754

**Authors:** Lauren E. Wilbanks, Elliot N. Brajkovich, Leo Baumgart, Yu Zhang, Nicolas Grosjean, Ian Blaby, Elizabeth I. Parkinson

## Abstract

Natural products (NPs) are a rich source of therapeutic and agricultural compounds. Unfortunately, many promising metabolites are not expressed under standard laboratory conditions. Deepening our understanding of the regulatory networks governing NP biosynthetic genes is essential for unlocking this hidden chemical diversity. Cluster-situated regulators (CSRs) are transcription factors involved in the regulation of NPs, but their full regulatory range has remained elusive due to limited genome-wide data. Using DNA Affinity Purification Sequencing (DAP-seq), we defined the predicted regulons for 84 CSR homologs across 78 *Streptomyces* strains. CSRs in this cohort exerted influence across multiple cellular processes, with particularly strong impacts on other transcription factors throughout the genome. Approximately 30% of predicted NP biosynthetic gene clusters (BGCs) contained CSR-regulated genes. In strains encoding multiple CSR homologs, we observed substantial overlap in BGC regulation. Together, these results greatly expand the genomic landscape of CSR activity and provide a foundation for improved bioinformatic strategies to predict and interpret regulatory control of NP biosynthesis.

## Background and Introduction

Bacteria employ many regulatory strategies to control transcriptional response to changes in their environment^1–5^. A major tactic is the use of transcription factors to regulate gene expression^6–8^. Transcription factors can function as either transcriptional activators, which bind to DNA and recruit RNA polymerase to a gene’s promoter, or repressors, which inhibit RNA polymerase binding to DNA^7,9–11^. To coordinate responses to external stimuli, groups of genes are often activated or repressed in a population dependent manner. Frequently, these systems are regulated by quorum sensing, which depends on small, membrane diffusible molecules. These quorum sensing molecules are made initially at low levels^1,2,4^. When the population density reaches a threshold, the increased concentration of quorum sensing molecules triggers gene activation, typically via a direct interaction with a transcription factor^2,12,13^. This tactic is employed by bacteria to regulate processes as diverse as luminescence, virulence, conjugation, and secondary metabolites^14–17^. Within the *Streptomyces* genus, quorum-sensing molecules called autoregulators (ARs) are frequently employed to regulate the production of Natural Products (NPs), secondary metabolites with great therapeutic and agricultural relevance^17– 23^. The enzymes that synthesize NPs are often encoded in genetic neighborhoods called biosynthetic gene clusters (BGCs). NP production demands the coordination of many processes, including production of precursors, transcription of NP biosynthetic enzymes, and regulation of proteins involved in resistance and export of the NP^6,24–27^. Additionally, for the NP to be synthesized in effective concentrations, its production must be coordinated throughout the bacterial population. Therefore, NP production necessitates tight regulation of numerous genes. Such regulation is often through repressors found within or near the NP gene cluster, named cluster-situated regulators (CSRs), and their associated ARs^28^. Modern bioinformatic analyses of *Streptomyces* genomes indicate there exists a vast repository of NPs not expressed under standard laboratory conditions, and extensive research has been performed attempting to elicit production of these silent or cryptic NP^29–35^. Recently, CSRs have emerged as regulators of interest in prioritizing NP BGCs for attempted NP elicitation^28,36–38,38–40^.

The majority of research on CSRs has focused on their roles as regulators of NP production, though in a handful of cases they have been found to regulate primary metabolism and sporulation. ^41–48^ Previous studies identified novel CSRs by their homology to known repressors, demonstrated their regulation of a proximal NP BGC, and proposed a consensus transcription factor binding site (TFBS) motif based on the binding sites for neighboring genes^45,46,49–55^. These investigations also proposed potential AR structures based on AR biosynthesis gene homologs within the CSR’s genomic neighborhood^56–59^. While this strategy has produced evidence for many CSR homologs, there remain critical gaps in our understanding of their full regulons, including the total number of NP BGCs they influence. Few genome-wide studies of CSR binding have been performed, with previous studies utilizing low-throughput methods, such as DNA footprinting analyses or electrophoretic mobility shift assays^46,50^. CHIP-seq provides for genome-wide TFBS identification, but would be challenging due to the need for numerous CSR-specific antibodies^60,61^. DNA Affinity Purification Sequencing (DAP-seq) is a low cost, high throughput alternative to CHIP-seq.^62^ Specifically, DAP-seq involves incubating a tagged transcription factor with isolated genomic DNA, with bound DNA being identified via genomic sequencing. Ultimately, this analysis enables generation of DNA-binding motifs for the transcription factor.^63^ This work details the analysis of *Streptomyces* CSRs from 78 strains, including the examination of the functions of the regulated genes. We particularly focused on regulated genes involved in NP and AR production. Understanding these essential CSR regulatory systems will yield insight into *Streptomyces* NP production, which will aid in the prioritization of novel NP discovery.

## Results and Discussion

### DAP-seq Experimentation and Predicted CSR Regulons

CSRs in this study belong to the TetR-like family of repressors, which possess a conserved N-terminus DNA binding domain and C-terminus ligand binding domain^66–69^. Two homodimers of CSR proteins bind to the TFBS within gene promoter regions (**Figure 3A**). Based on published DNA-CSR complexes of AvaR1 from *S. avermitilis* and MmfR from *S. coelicolor*, TFBSs for CSRs are broadly comprised of two core binding motifs (e.g. ACCG/CGGT), separated by a 6 bp spacer (**Figure 3B**). Of these core motifs, the AC/GT is typically highly conserved, however, these core binding motifs frequently exhibit imperfect sequence symmetry (**Figure 3B**)^66^. To identify TetR-like repressor homologs, a sequence similarity network (SSN) was built using the EFI-EST tool (**Supplementary Dataset 1**)^64^. ScbR, a TetR-like repressor from *Streptomyces coelicolor* (Accession WP_003972660), was used as the input sequence. The resultant SSN contained several well-known TetR-like repressors, along with over 900 additional CSR homologs (**Figure 1**). 92 CSRs were selected from different SSN groups to determine their TFBS motifs using DAP-seq (**Figure 1**). Typically, DAP-seq is performed with genomic DNA from the corresponding organism, however, many of the *Streptomyces* strains within the SSN were not available^62^. For this reason, genomic DNA was isolated from 11 diverse *Streptomyces* strains for use as the DAP-seq template. All CSRs were screened against extracted DNA from these 11 strains to ascertain if DAP-seq can be used to determine a CSRs’ regulon even when the strain is not available. For each CSR, binding sites from the assays against each of the 11 *Streptomyces* DNA isolates were compiled and MEME was used to generate binding motifs (**Figure** ). Out of the initial 92 CSRs, 84 CSRs produced binding results sufficient for bioinformatic analyses. The canonical structure of CSR TFBS motifs was used to guide the MEME parameters for motif generation (**Figure 3B, Supplementary Figure** ). Initially, an 18-20 base motif was generated followed by a smaller motif window to identify the extent of the conserved regions (**Supplementary Dataset** and **Supplementary Figure** ).

**Figure 1.**
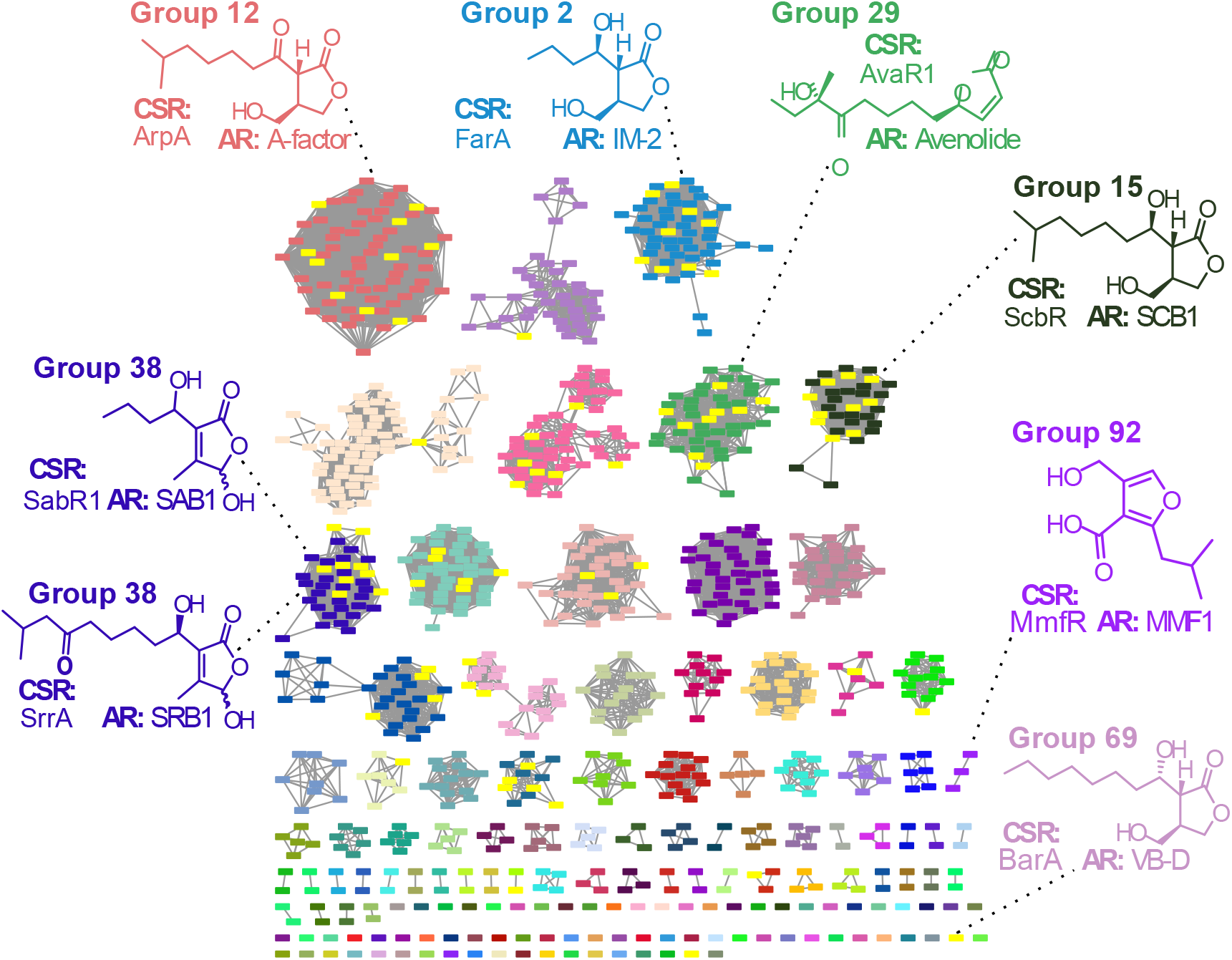
Sequence Similarity Network of *Streptomyces* CSRs. Each cluster has a unique color. CSRs evaluated by DAP-seq are yellow. The known CSRs and cognate ARs are indicated along with their SSN groups. Full legend is in **Supplementary Figure 1**.

**Figure 2.**
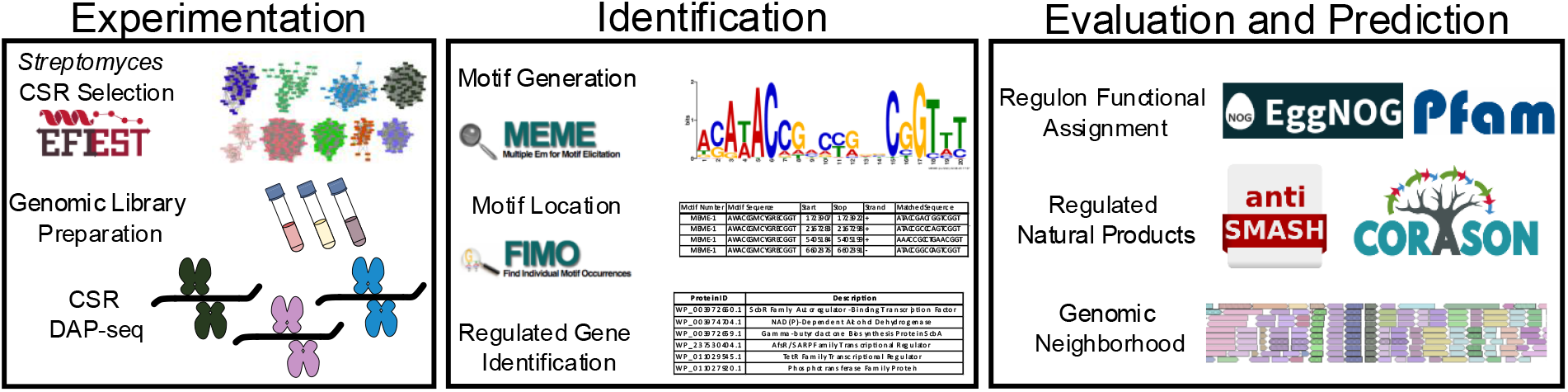
Project overview.

**Figure 3.**
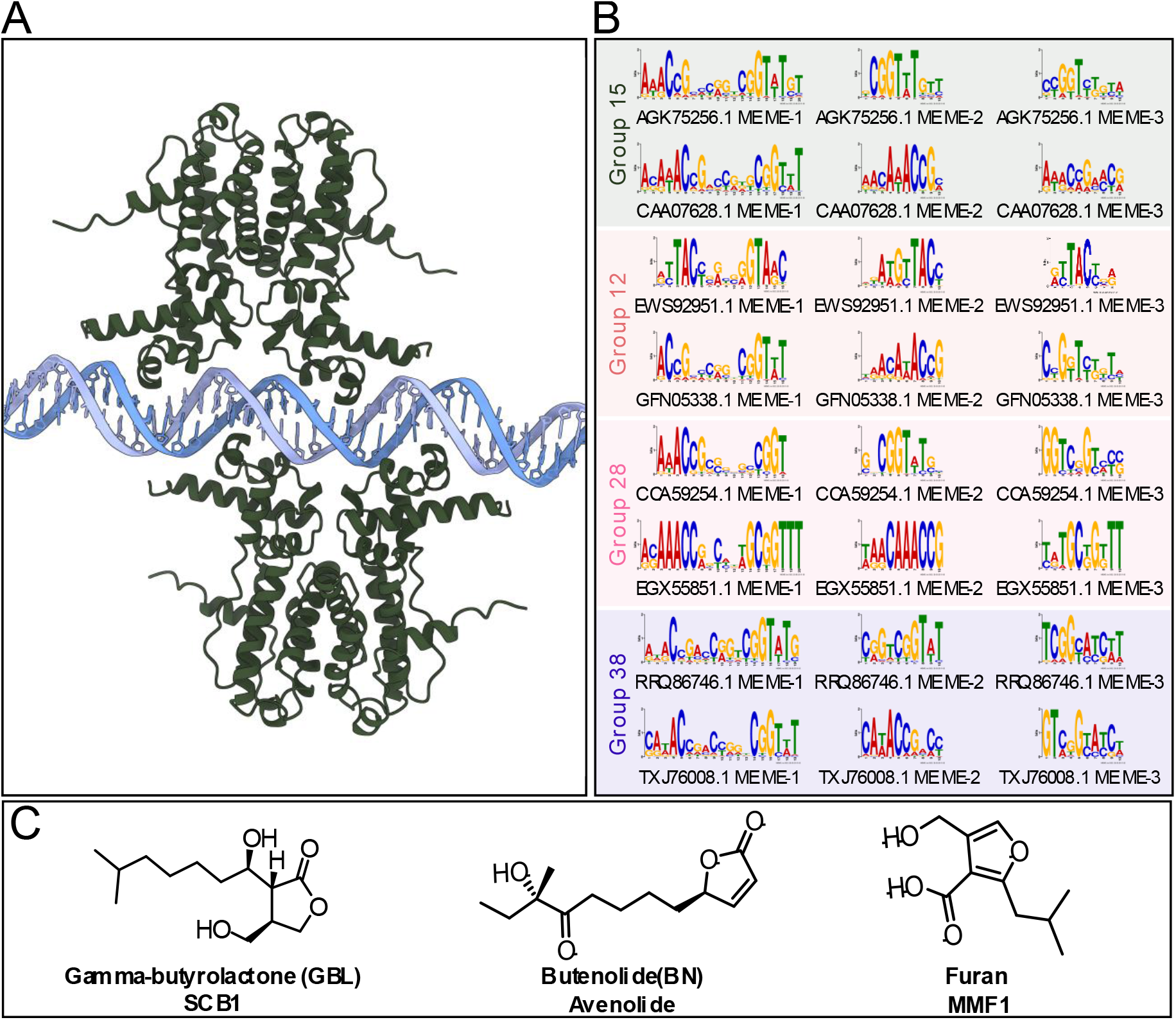
Structure of CSRs and cognate autoregulators (ARs). A) AlphaFold3 Model of two CSR dimers (ScbR) bound to its TFBS. B) MEME outputs of CSR from four SSN Groups C) Examples of different classes of *Streptomyces* NP ARs.

Once the motifs were determined, FIMO (Find Individual Motif Occurrences) was used to identify motif locations within each CSR genome (**Supplementary Dataset 3**). These binding locations were reduced to only include sites within gene promoter regions. Initial regulated gene hits were narrowed based on number of binding sites within the gene’s promoter to afford increasingly selective libraries of regulated genes (**Supplementary Dataset 4**). For the most stringently regulated gene library, eggNOG and Pfam were used to identify functions of regulated genes^73,74^. The overwhelming majority of predicted regulated genes were involved with transcriptional regulation or signal transduction; this was consistent in every CSR cluster (**Figure 4**). The numerous families of transcription factors that these CSRs regulate speak to their essential role within a network of regulators that direct genetic flux in response to external signals. Specific regulated families include stress response and antibiotic resistance regulators (MarR/MerR/TetR *n* = 284), carbon, nitrogen and amino acid regulators (PucR/LacI/AsnC, *n* = 80), and global regulators of secondary metabolism/morphogenesis (AfsR/SARP/LysR/IclR/AraC, *n* = 59)^6^. Other common targets of CSR regulation were purported AR biosynthesis genes, including annotated genes such as ScbA-like biosynthesis enzymes, NAD(P)/NAD(P)H-oxidoreductases, or acyl CoA dehydrogenases^53,66,75^.

**Figure 4.**
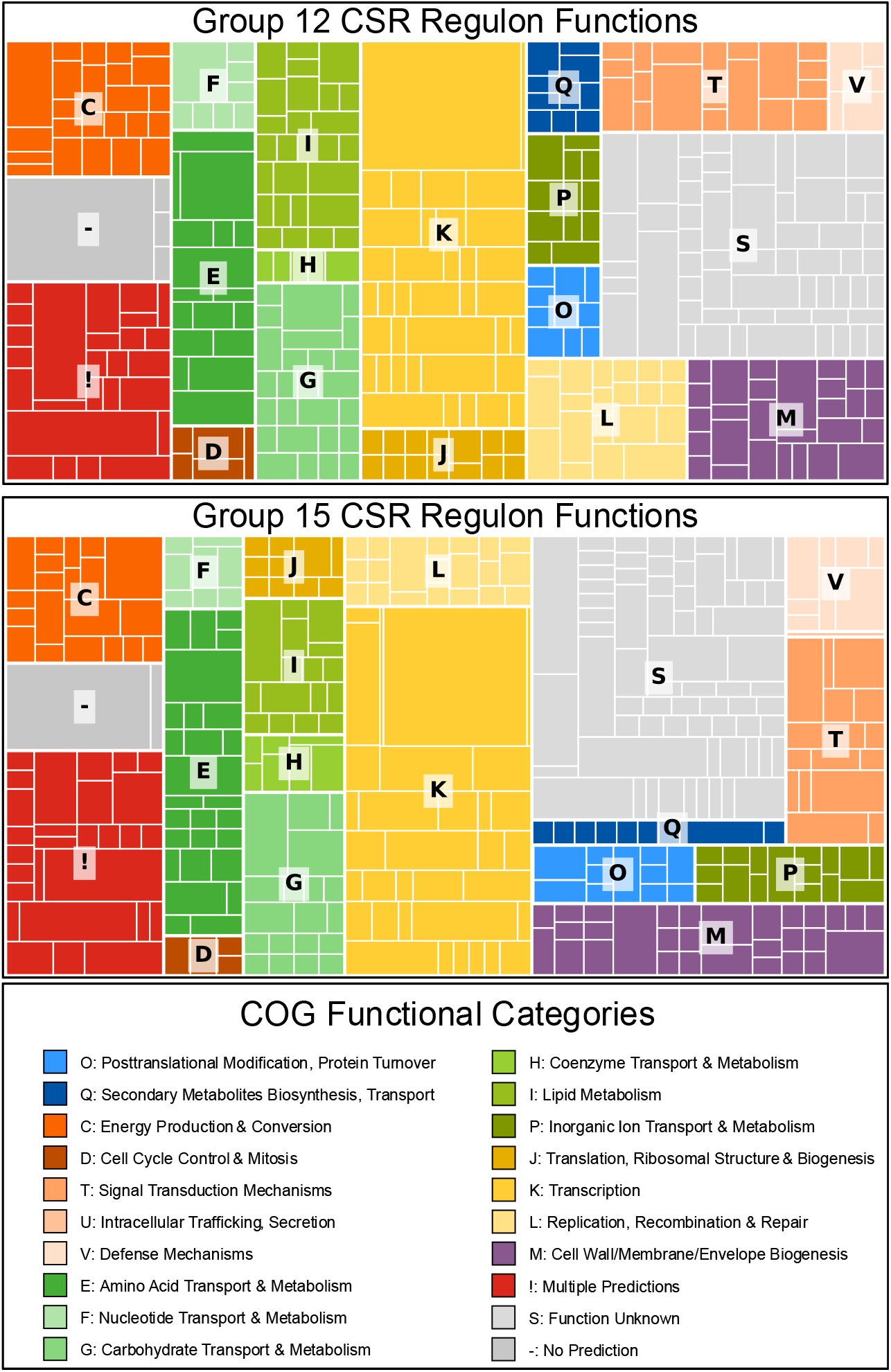
Functional Analysis of Genes in CSR Predicted Regulons. Treemap with COG functional categories for CSR SSN Groups 12 and 13 are presented. Subsections within treemap indicate unique Pfam identifiers.

Another top target included other TetR homologs, annotated in RefSeq as ‘ScbR family autoregulator-binding transcription factor’. A survey of the 78 genomes indicated each strain had on average 4 of these homologs with the analyzed CSRs regulating an average of one additional homolog (**Supplementary Dataset** ). These homologs were frequently present within the same genomic neighborhood. As predicted, there were numerous functional annotations for secondary metabolite production (**Figure 4**). Interestingly, the analysis also indicated an abundance of regulated genes that were homologous to primary metabolic enzymes. Further investigation is needed to determine if these are involved in primary metabolism or if these genes have been evolutionarily repurposed as NP biosynthesis genes^76^. Overall, these findings reinforce well known roles of CSRs and serve to validate the bioinformatic pipeline for regulated gene identification. From this broad view of CSR regulons, it is evident that they function as regulators of regulators, leading signaling cascades and gene fluxes when benchmarks indicate the cell is ready to activate secondary morphogenesis and NP expression.

### CSR-Regulated Genes within NP BGCs

We sought to utilize our bioinformatic results to identify NP BGCs that are regulated by CSRs. The 78 genomes of origin of the CSRs were processed using antiSMASH to identify NP BGCs. 34 of the genomes were chromosome-level assemblies, and 44 genomes were scaffold-level assemblies. The regulated genes within the ‘Top Hits’ library were searched against the BGCs. A NP BGC was considered for further analysis if it had at least one gene from the ‘Top Hits’ regulated gene library within its BGC. The CSRs examined in this study exercise wide influence on numerous NP BGCs within their genomes. Each CSR was predicted to regulate multiple NP BGCs from all NP classes. CSRs regulated at least one gene in 11%-59% of predicted NP BGCs, with an average of 30% of identified NP BGC being regulated by a CSR (**Supplementary Dataset** ). Of the approximately 850 BGCs with gene(s) regulated by CSRs, only approximately 370 BGCs shared >60% similarity with a known cluster within the MiBIG database.^77^ nown CSR-regulated NPs correctly called by the analysis include ScbR/CAA07628.1 regulating coelimycin P1, SrrA/BAC76540.2 regulating lankamycin, and BarA/BAA06981.1 regulating virginiamycin S1^44,47,78,79^. Functional assignment of NP BGC regulated genes was performed using eggNOG^73^. Predictably, the most prevalent gene function was transcription (∼500 assignments) (**Supplementary Table** ). Amino acid and lipid transport and metabolism, alongside ‘secondary metabolites biosynthesis, transport, and catabolism’ (each have ∼150 assignments) were also commonly observed.

*Streptomyces microflavus* (GCF_045039785.1) emerged as a genome of special interest, as two CSRs from the DAP-seq cohort are present within this genome. With this data, we were able to determine the interaction of the CSR regulons, with particular interest in the NP BGCs that they regulate (**Figure** ). The CSRs come from different SSN groups QKW44091.1 belongs to Group 12 and AGK75256.1 to Group 15. Individually, QKW44091.1 and AGK75256 regulate genes within ∼30% of the NP BGCs within the strain. For different NP BGCs in *S. microflavus*, these CSRs were found to often regulate identical genes within the same cluster or different genes within a cluster, resulting in them together regulating genes from 40% of the NP BGCs in *S. microflavus* (**Figure 5**). Interestingly, both QKW44091.1 and AGK75256.1 regulate the same genes in the butyrolactone Cluster 3: a *scbA* homolog, AGK75256.1, which likely is part of an AR BGC due to its similarity to known AR biosynthesis enzymes *afsA and bprA* (**Figure 6C**). The known CSR from Group 12 (ArpA) responds to A-factor while the known CSRs from Group 15 (ScbR) responds to SCB. Interestingly, A-factor and the SCBs differ only in the oxidation state of the C1’ position (**Figure 1**). Previously, we have found that there can be crosstalk between these CSRs (*i.e*. A-factor can derepress ScbR, and the SCB derivative SCB1 can derepress ArpA), although with markedly lower affinity compared to that of the native ligand^80–82^. Thus, it is possible that QKW44091.1 and AGK75256.1 may crosstalk within *S. microflavus*. Given this potential crosstalk between both TFBS and AR, it appears that the regulation by QKW44091.1 and AGK75256.1 in *S. microflavus* are interdependent. Similar coregulation also exists in *S. microflavus* (GCF_036227005.1), and *Streptomyces* sp. S8 (GCF_002094995.1), with both having CSRs from groups 12 and 15 that show overlapping regulation of NP BGC genes. *Streptomyces* sp. MMG1533 and *Streptomyces* sp. WM6378 also have overlapping regulation of the tested CSRs, but their CSRs belong to Groups other than 12 and 15. Taken all together, this data proposes an expansive and interconnected NP regulon for CSR homologs within a genome.

**Figure 5.**
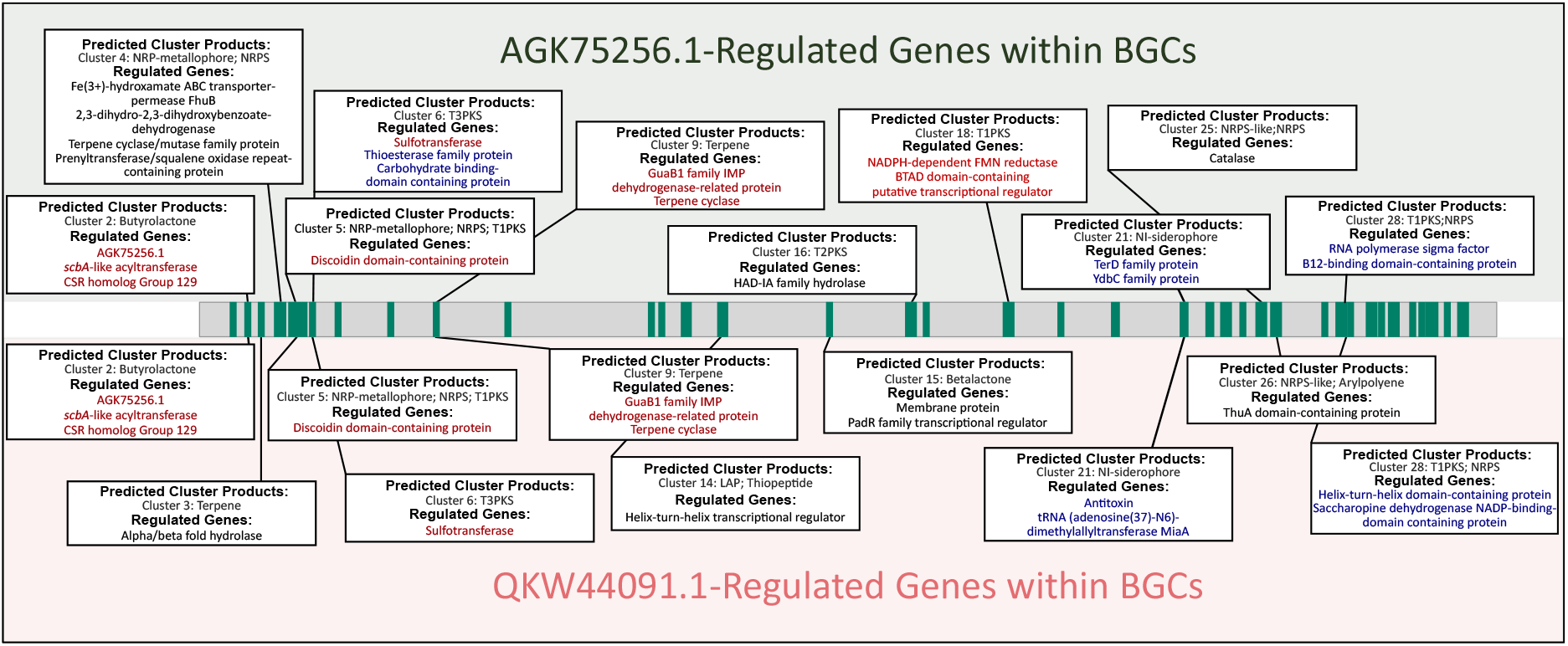
Interacting S regul ns in *Streptomyces microflavus*. Green bars are predicted NP BGCs in the *S. microflavus* genome, top boxes indicate regulated BGCs of AG 75256.1, lower boxes indicate regulated BGCs of W44091.1. Red annotations are for shared regulated genes. Blue annotations are for genes where the CSR regulate members of the same BGC but not the same gene. Black annotations are genes within a BGC regulated by only one CSR.

**Figure 6.**
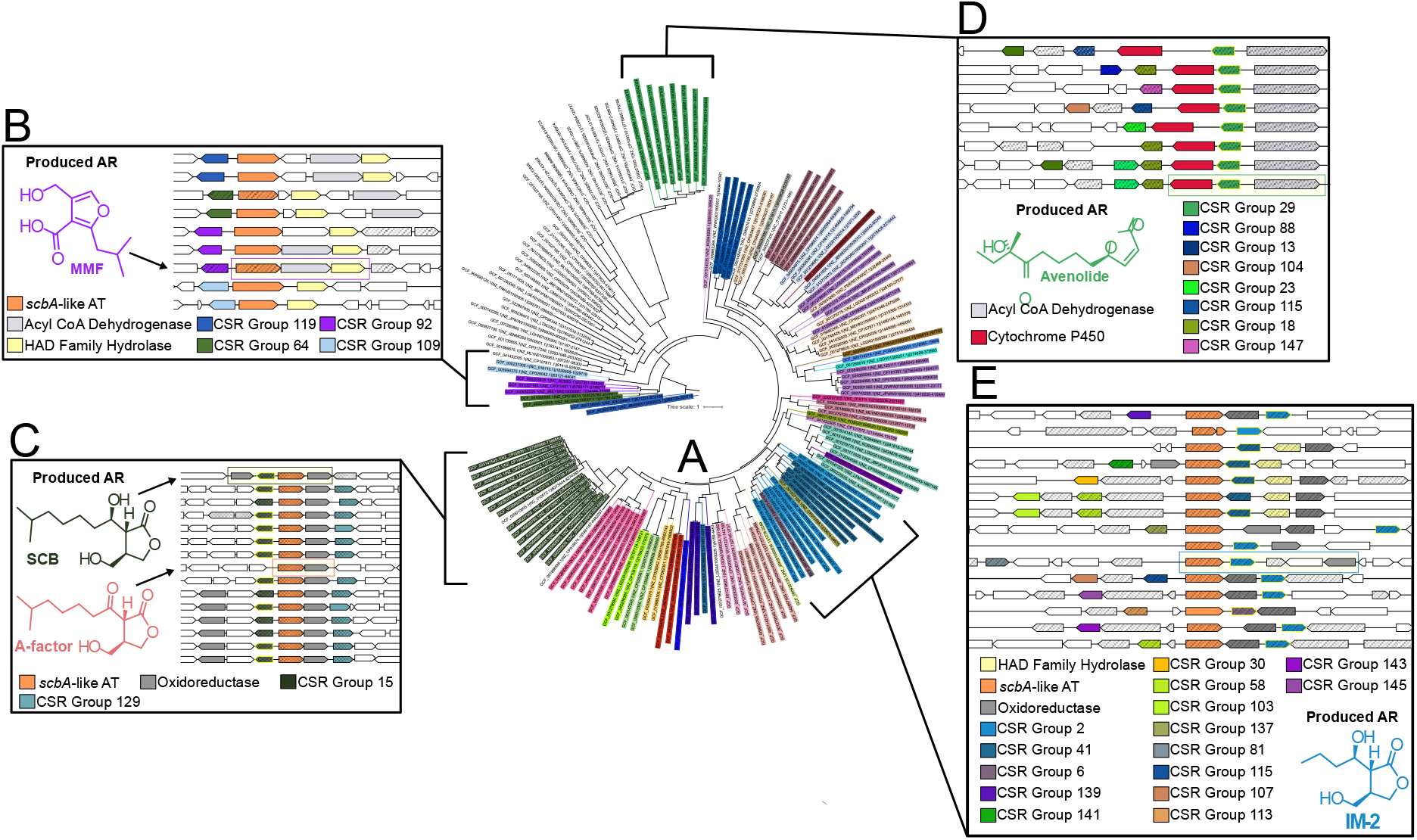
Phylogenetic tree of AR primary synthesis enzymes with associated genomic neighborhoods. Phylogenetic tree of AR primary biosynthetic enzymes is in the center (A), with the indicated genomic neighborhood of the primary enzyme (B-E). AR biosynthetic enzymes including *scbA*-like (orange) or *aco*-like (light grey) were used as anchors for constructing the tree. Tree labels are colored by the SSN group of the CSR present in the AR BGC; clusters with multiple CSR homologs are colored based on the CSR most proximal to the primary enzyme. Uncolored labels do not have a CSR homolog in their BGC. Genes for AR biosynthetic enzymes are highlighted, with known AR structures showed. AR biosynthetic genes and CSR groups are color coded as indicated; regulated genes are striped; DAP-seq CSRs are outlined in yellow.

### Genomic Neighborhood of Autoregulator BGC with CSR homologs

nowledge of the ARs that bind CSRs is essential for fully understanding these regulatory networks. The structure of the CSR ligand binding domain determines the type of AR that it binds. nown CSR AR classes include butenolides (BNs), furans, and gamma-butyrolactones (GBLs) (**Figure 1** and **3** ). The first step in GBL biosynthesis is a condensation of a β-keto ester and DHAP, performed by a homolog of the *S. coelicolor* acyltransferase *scbA*,^53,56–58,83^ with further tailoring of the molecules being performed by various oxidoreductases^53,75^. Biosynthesis of furans and the SAB and SRB butenolides also begin with an *scbA* homolog, with the products being being subjected to different oxidoreductases to afford their diverse final structures (**Figure 1** and **3** )^44,45,54,55^. The biosynthesis of avenolide, the butenolide AR from *S. avermitilis*, is an outlier and is instead believed to be performed by an acyl-CoA dehydrogenase and a cytochrome P450 family protein^66^. Previous work indicates that AR biosynthetic genes and their cognate CSRs are typically encoded within close proximity, with the CSR and the initial biosynthesis enzyme sharing a bidirectional promoter^50,66,75,84^. This work also found that CSRs regulate both their own expression, as well as the initial biosynthesis enzyme. We sought to combine the CSR homology analysis provided by the SSN with the genomic neighborhoods of the AR BGC and data on CSR regulated genes to guide AR-CSR prediction.

AR BGCs called by antiSMASH (named as either ‘furans’ or ‘butyrolactones’) were analyzed alongside avenolide-type AR BGCs identified by CORASON. 76 of 78 (97%) of genomes examined have at least one AR BGC, with an average of two and a maximum of 4 per genome. 62 of 84 (74%) of the CSRs that were examined by DAP-seq reside within a predicted AR BGC. Of the 149 ‘butyrolactone’ BGCs called by antiSMASH, 73 were regulated by the CSRs (∼50%). Interestingly, ∼50 of the butyrolactone BGCs were not situated proximal to another called NP BGC. That is, the only NP produced in these clusters was the butyrolactone. Only 9 furan BGCs were predicted, and 2/9 of these furan BGC possessed genes that are regulated by a CSR. Currently, avenolide-type BN BGC are not called by antiSMASH. To ascertain the number of avenolide-type BN we used CORASON with our 78 *Streptomyces* genomes^85^. An *aco* (acyl CoA dehydrogenase) homolog was used as the anchor protein for CORASON analysis with our *Streptomyces* genomes, which resulted in 17 predicted avenolide BGCs. To observe the evolutionary similarities between the AR BGCs, a phylogenetic tree was built using the primary biosynthetic gene as an anchor, either *aco*-like for avenolide-type BN, or *scbA*-like for GBL, furan, and other BN-type ARs (**Figure A**). All CSR homologs present within the AR BGC were identified, as well as any CSR regulated genes. As expected, many of the AR BGCs within a clade possessed a CSR from the same SSN group. An evolutionary similarity also emerged between AR BGCs that did not possess a CSR homolog.

The furan-type AR MMF has been experimentally determined to be produced by biosynthetic enzymes on the SCP1 plasmid in *S. coelicolor*^54,55,67^. The acyltransferase MmfL performs the initial step in MMF synthesis, while MmfP performs a dephosphorylation and MmfH catalyzes a rearrangement to afford the final MMF derivative structures (**Figure B**). The Group 92 CSR MmfR is present immediately upstream of the biosynthetic genes. The Group 109 CSRs lack *mmfP*, however, it has been reported that MMF can be made in the absence of *mmfP*^55^. It is hypothesized that other homologs within the genome may perform these functions^55^. Based on the evolutionary closeness of the acyltransferases and presence of the other AR biosynthetic homologs, it is highly likely that CSRs present in SSN groups 64, 92, 119, and 109 are all regulated by MMF-type ARs. Although MmfR was not included in the DAP-seq experimental cohort, ScbR/CAA07628.1, also present in *S. coelicolor*, was found to regulate multiple genes on the SCP1 plasmid, including *mmfL*, as well as *mmfR*. During *in vitro* testing, SCB1 did not demonstrate derepression of MmfR, and MMF1 did not derepress ScbR^80^. However, it appears that ScbR, and thus by extension SCB1, can regulate MMF and MmfR.

In Figure 6C, the biosynthetic genes for the production of the ARs SCB (from *S. coelicolor*) and A-factor (from *S. griseus*) are highlighted. It is logical that the acyltransferase is highly homologous given the high structural similarity of these ligands. All AR BGCs also possess a homolog of the butenolide reductase (*brpA* in *S. griseus* and *scbC* in *S. coelicolor*) immediately downstream of the acyltransferase. The NADPH-dependent oxidoreductase (*scbB*) immediately upstream of the Group 15 CSR is only present in the *S. coelicolor* AR BGC. Several clusters possess an oxidoreductase further upstream, but these enzymes share low homology to *scbB*. The enzymes present in the AR BGCs suggest that most of the AR BGCs in this clade make the A-factor type GBL with an exocyclic ketone, and not the SCB type with an exocyclic hydroxyl. All AR genomic neighborhoods other than the A-factor and SCB BGC possess a Group 129 CSR homolog which are consistently regulated by the Group 15 CSR. Of note, no SSN Group 12 CSR is present within these AR BGC, despite A-factor being the established cognate AR of ArpA. *arpA* is a known regulator of *adpA*, itself a master regulator of secondary morphogenesis in *S. griseus. arpA* is located 4.9 MB upstream of the A-factor BGC. The lack of Group 2 homologs, together with the presence of Group 129 homologs, underscores that proximity to an AR BGC alone is insufficient for identifying AR–CSR cognate pairs.

The avenolide clade with Group 29 CSR homologs provides several interesting observations for the acquisition and evolution of CSRs and AR BGCs (**Figure D**). AvaR1, the established receptor of avenolide, is the canonical CSR of Group 29. AvaR1 was submitted as part of the DAP-seq cohort but did not produce usable results. All avenolide-type ARs possess both the purported AR biosynthesis genes acyl CoA dehydrogenase and cytochrome P450. Additionally, the acyl CoA dehydrogenase and Group 29 homolog share a promoter region, and the Group 29 CSRs are predicted to regulate the acyl CoA dehydrogenase^66^. This high level of conservation is interesting in that the CSR from *S. lunaelactis* (A Z77953.1) is on a plasmid, where the other members of this cluster are genomic. CSRs from this cluster also regulate a number of other regulators within their genomic neighborhood, including AfsR/SARP family, TetR/AcrR family, and CSR homologs from a number of different SSN Groups. The CSR from *S. pristinaespiralis* (AA 07686.1) is present twice in the genome, at ∼412 B and another paralog at ∼8.1 MB, near the beginning and the end of the chromosome, which is commonly where NP BGCs reside in *Streptomyces*. It is unclear if these paralogs are the result of a double insertion of a linear plasmid carrying an AR-regulated BGC, if one end of the *Streptomyces* chromosome broke off and recombined, or some other gene duplication mechanism occurred^86–88^. One other CSR we examined, RWZ75665.1 from *S. albidoflavus*, also exhibited a similar pattern of two paralogs with identical genomic neighborhoods near the ends of the chromosome.

The known CSR of SSN group 2 is FarA, which is regulated by the gamma-butyrolactone IM-2^48,50^. The AR BGC of IM-2 is identical to the highlighted BGC, possessing an *scbA*-like acyltransferase, a GNAT family N-acetyltransferase (regulated protein proximal to the CSR), and a NAD(P)H-binding protein (**Figure E**)^83^. The key difference in this ligand from other GBL-types is its short acyl chain. While A-factor and SCB1 have 7 carbon chains, IM-2 has a 4 carbon chain (**Figure 1**). Empirically, IM-2 does not de-repress ScbR or ArpA; likewise SCB and A-factor did not activate FarA^80^. We have hypothesized that the short sidechain lends, as larger GBL-types A-factor and SCB1 likely cannot bind FarA^80^. This ligand diversity would explain the distance between the acyltransferase of SCB/A-factor and IM-2, as the IM-2 *scbA*-type acyltransferase is likely adapted to work on a substrate with a short sidechain. Group 41 CSRs are also present in this clade, but all of these AR neighborhoods have a HAD hydrolase. Although there is similarity between the *scbA*-type acyltransferases of Group 41 and Group 2, the CSR belonging to different SSN groups and the presence of the HAD family hydrolase is highly suggestive of a different final structure. While Group 2 and Group 41 CSRs appear consistent throughout the observed AR BGCs, the other CSR homologs represent diverse SSN groups. Further investigation will reveal the extent of structural diversity of their cognate ARs.

## Conclusions

A plethora of novel NPs reside within *Streptomyces* but are unexpressed under standard laboratory conditions^18,29,31,89^. Elucidating CSRs that regulate NP production can aid in selection of NP BGC candidates for NP elicitation^37,38^. To manipulate CSR regulation, researchers must identify their TFBS and thus their regulated genes, as well as establish an AR ligand for potential induction. Starting with DAP-seq binding data, our bioinformatic pipeline can determine CSR TFBS motifs and locate predicted regulated genes. These regulons can be mined to inform on the roles of CSRs in regulating various processes, including appraising their participation in regulation of NP production. The regulons the pipeline generated confirmed the role of CSRs as regulators of other regulators, along with NP BGCs. While further experimentation will be necessary to confirm regulated genes, this work provides a resourse to prioritize gene validation based on binding data. A SSN provides examples of structurally similar CSRs, and can link CSRs under investigation to CSR homologs with existing literature precedent to provide information on cognate ARs^64^. The genomic neighborhood of a CSR may also provide additional aid when linking CSRs to their ARs, as homologs to known biosynthetic genes proximal to the CSR can provide a strong prediction of the CSR’s cognate AR^84^. Looking at regulated genes within NP BGCs, we confirmed known CSR-regulated NPs and identified CSRs that participate in regulating on average a third of predicted NP BGCs within the genome. For genomes where we possessed regulon data on two CSRs, we observed coordination between the CSRs, both coregulation of individual genes or regulation of different genes in the same BGC. In total, this work provides a bioinformatics-guided structure for exploring CSR regulons, with the ultimate goal of identifying NP BGCs that are candidates for elicitation via induction with AR.

CSR/AR systems are a widespread mechanism for the coordination of primary and secondary metabolism for NP production in *Streptomyces*. The NCBI database has ∼2300 proteins similar to the CSRs assayed in this research; these regulators are present in ∼1000 unique *Streptomyces* genomes^59,90,91^. These regulators appear essential for *Streptomyces* to preserve an extensive collection of NP BGCs. As transcriptional repressors, the CSRs maintain an ‘OFF state’ in which the expression of genes they regulate is silent. The high similarity in TFBS indicates that there is crosstalk in regulation between CSRs, with a single gene potentially being the target of multiple CSRs. Crosstalk between CSRs and the operators of other CSRs has been empirically demonstrated^80^. While the DNA binding domains afford TFBS promiscuity, the ligand binding domain is more selective. ARs of different classes show low to no ability to activate non-cognate CSRs^80^. CSRs that respond to different ARs may regulate the same genes within a BGC, or each CSR may regulate unique genes within one BGC. If multiple CSRs that bind different classes of ARs regulate a single BGC, all cognate ARs must be present in the proper concentrations to completely derepress the CSRs and produce the NP. Investigations into *Streptomyces* CSRs cannot solely look at individual CSRs but must examine the broader context in which they interoperate. This sophisticated regulatory bureaucracy appears to be essential for governing NP production, sporulation, and general respond to external threats.

### Experimental Section

Procedures can be found in the Supplementary Information document.

## Associated Content

### Data Availability Statement

Supporting datasets are available at the Purdue University Research Repository (PURR, add DOI once published).

### Supporting Information

The Supporting Information is available free of charge.

Supporting information, including methods, supplementary tables, and supplementary figures.

## Acknowledgements

The work (proposal: https://doi.org/10.46936/10.25585/60008554) conducted by the U.S. Department of Energy Joint Genome Institute (https://ror.org/04xm1d337), a DOE Office of Science User Facility, is supported by the Office of Science of the U.S. Department of Energy operated under Contract No. E-AC02-05CH11231. This work was funded by an NSF CAREER Award to E.I.P. (CHE 223689). L.E.W. was supported by a Purdue Research Foundation Ross-Lynn Grant. This work was supported in part by the Research Instrumentation Center in the Department of Chemistry at Purdue University. The authors acknowledge the support from the Purdue Center for Cancer Research, NIH grant P30 CA023168, with particular thanks to the C3B-Collaborative Core for Cancer Bioinformatics.

