## Supplementary Information and Methods for "DAP-seq Reveals Cluster-Situated Regulator Control of Numerous *Streptomyces* Natural Product Biosynthetic Genes"

#### Methods and Materials

*TF selection and DAP-seq experimentation*

EFI-EST tool was employed using the amino acid sequence from the CSR ScbR from *Streptomyces coelicolor* as the template[1]. The sequence similarity network (SSN) was generated using a 65% sequence identity threshold and a 70% alignment score cutoff (65i/70c). The resulting sequence similarity network was visualized in Cytoscape v3.10.2[2]. Initially, 92 CSR sequences were identified from a number of different clusters generated by the SSN. The coding sequences for these CSR were provided to the Joint Genome Institute (JGI), along with 11 *Streptomyces* genomes as templates for the DAP-seq analysis (**MethodsTable 1**). *Streptomyces* genomes were isolated using the Qiagen DNeasy PowerLyzer PowerSoil Kit in accordance with a published protocol[3]. DAP seq was performed as previously published, and all 92 CSR were assayed against all 11 genomes in duplicate[4]. Each transcription factor sequence was iteratively refactored in BOOST using an *E. coli* codon frequency table until all synthesis violations were resolved. Codon refactoring employed the "balanced" strategy, which aligns codon usage with frequencies in the *E. coli* genome. Thirty–base pair DNA linkers were appended to the 5′ and 3′ ends of each sequence to enable scarless, in-frame assembly downstream of a HALO tag in pIX-Halo_PaqCI, which had been linearized by PaqCI digestion. Linear synthetic DNA fragments (Twist Bioscience, CA, USA) were PCR-amplified and assembled into the linearized vector using the NEBuilder Hi-Fi DNA Assembly kit (New England BioLabs). All constructs were sequence-verified on the Pacific Biosciences Sequel IIe platform and analyzed with custom pipelines[5]. From this cohort, 84 CSRs were identified for further investigation based on sufficient binding activity during. CSRs were dropped from further analysis if no binding occurred during the assay or if the identified TFBS had a fold change of <15.

*DNA-ScbR model*

DNA-ScbR model was generated using AlphaFold3[6]. The TFBS sequence used was 5' - GGACAAGCGCCATCGGAACCGGCAATGCGGTTTGTTCGA-3'; ScbR/CAA07628.1 amino acid sequence is listed in **Supplementary Dataset** 1.

*TFBS motif generation and location, analysis of CSR regulated genes*

Raw DAP-seq peak sequences from all runs were compiled and curated for each CSR. Peak sequences were enriched or removed based on fold-change. Sequences were removed from analysis if their fold change was <15. Additional entries of high fold change sequences were added, multiplied based on fold change. Curated peak sequences were processed using MEMEsuite 'MEME' to identify binding motifs[7]. The unique structure of the *Streptomyces* CSR TFBS necessitated two independent runs to produce reliable motifs for each CSR. The first motif (i.e. motif 1) was identified by adjusting the minimum/maximum motif width parameters such that the output conformed to the conserved TFBS structure exhibited by a number of known CSRs (e.g. ACNNNNNNNNNNGT)[8]. Motifs 2 and 3 were generated by adjusting the parameters to generate two pseudo-palindromic motifs (e.g. AAAC / GTTTT). For each CSR, these three motifs were combined into a single input, then scanned against the CSR genome of origin using MEMEsuite FIMO to identify genomic locations that matched the motifs. These TFBS were processed to identify exclusively motifs that were present in annotated gene promoter regions. Additional sorting was performed based on number and location of motif binding, generating three libraries of regulated genes with increased stringency: **All**: Gene hits that have motif 1, 2, or 3 present in the promoter region; **Top Hits**: Hits from 'All' that have at least one 'motif 1' present in the promoter region; **Curated**: Hits from ‘Top Hits’ that have >= 2 motifs present in the gene promoter, at least one ‘motif 1’ and another incidence of motif 1, 2, or 3. The eggNOG database v5.0 was used to assign COG terms to regulated genes. The ‘Curated’ regulated gene library for each CSR SSN Group were combined, then used as the basis to generate treemaps of the COG terms[9,10].

*NP regulation and AR BGC analysis*

For each genome, we sought to identify NP BGCs that possessed genes predicted to be under regulation of a CSR, thus identifying NP production that could be regulated by our library of CSRs. First, antiSMASH v7.1.0 was run on all 78 unique genomes with detection strictness ‘relaxed’ to identify predicted BGCs. Additional analyses were performed to attempt to identify potential butenolide BGC. CORASON was run using the anchor proteins the acyl-CoA dehydrogenase from *S. lunaelactis* (WP_108155242.1), as well as the ScbA-like acyltransferases WP_011113166.1 from *S. rochei* and WP_229896777.1 from *S. eurythermus*. CORASON was run using default settings, with a BLASTp e-value cutoff of $1 \times 10^{-15}$ and a ±10 gene window[11]. Redundant BGCs were removed to provide the total predicted AR BGC library. Anchor proteins were identified as either acyl CoA dehydrogenase for avenolide-type butenolide ARs, or ScbA-like acyltransferases for GBL-type ARs. Anchor proteins were aligned using MAFFT v7.505 using the local pairwise alignment + iterative refinement method[12]. Alignments were performed with default gap penalties under the BLOSUM62 substitution matrix. ModelFinder Plus (MFP) was used to select the best-fitting amino acid substitution model[13]. The final maximum-likelihood tree was generated using IQ-TREE2[14]. The phylogenetic tree was visualized using ITOL, gene clusters were visualized using DNA Features Viewer[15,16]. After all AR biosynthesis genes were identified, genes from the ‘All’ library were searched against the identified BGC regions to determine NP BGC that were regulated by the CSR.

**Methods Table 1**. Strains Used for DAP-seq DNA Template

| Strain Name | Source |
|---|---|
| *Streptomyces avermitilis* ATCC 31267 | ATCC |
| *Streptomyces rochei* D21E05 | Ju (2015)[17] |
| *Streptomyces lividans* 66 | John Innes Centre culture collection |
| *Streptomyces gardneri* ATCC 15439 | ATCC |
| *Streptomyces antibioticus* DSM 41481 | DSM |
| *Streptomyces griseofuscus* DSM 40191 | DSM |
| *Streptomyces clavuligerus* NRRL 3585 | NRRL |
| *Streptomyces sp.* NRRL B-3648 | NRRL |
| *Streptomyces sp.* NRRL S-4 | NRRL |
| *Streptomyces sp.* MMG1533 | Ju (2015)[17] |
| *Streptomyces alboflavus* NRRL B-2373 | NRRL |

### Supplementary Information

**Supplementary Table** 1. COG Assignments for Regulated Genes with NP BGCs

| COG Category | Total Assignments |
|---|---|
| Transcription | 510 |
| Amino Acid Transport/Metabolism | 148 |
| Carbohydrate Transport/Metabolism | 130 |
| Secondary Metabolites Biosynthesis/Transport/Catabolism | 130 |
| Signal Transduction Mechanisms | 123 |
| Lipid Transport/Metabolism | 114 |
| Energy Production/Conversion | 81 |
| Cell Wall/Membrane/Envelope Biogenesis | 74 |
| Inorganic Ion Transport/Metabolism | 72 |
| Coenzyme Transport/Metabolism | 53 |
| Replication/Recombination/Repair | 50 |
| Translation/Ribosomal Structure/Biogenesis | 39 |
| Nucleotide Transport/Metabolism | 33 |
| Defense Mechanisms | 24 |
| Posttranslational Modification/Protein Turnover/Chaperones | 21 |
| Cell Cycle Control/Division | 9 |
| Intracellular Trafficking/Secretion/Vesicular Transport | 5 |
| Cytoskeleton | 1 |
| Function Unknown | 391 |

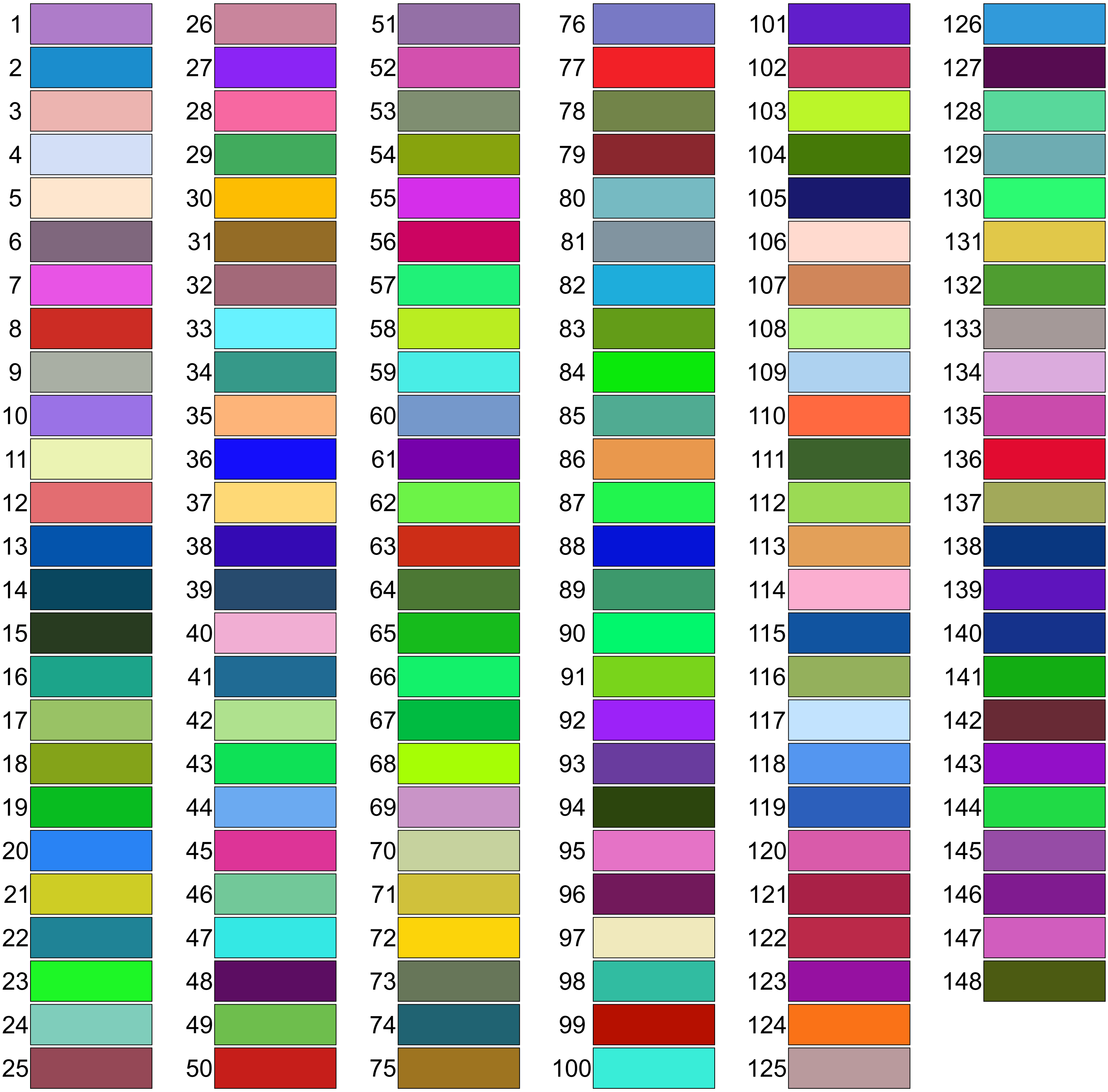

**Supplementary Figure 1.** Full legend of Sequence Similarity Network groups. SSN group number is next to figure color.

#### Group 1

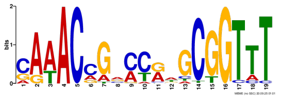
KOU47561.1 MEME-1

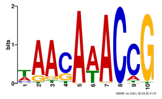
KOU47561.1 MEME-2

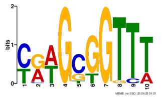
KOU47561.1 MEME-3

#### Group 2

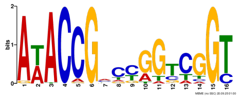
ANJ11867.1 MEME-1

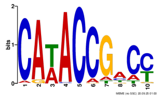
ANJ11867.1 MEME-2

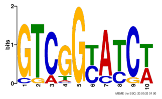
ANJ11867.1 MEME-3

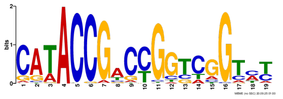
BAG74716.1 MEME-1

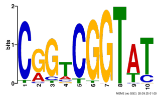
BAG74716.1 MEME-2

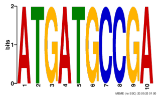
BAG74716.1 MEME-3

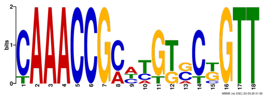
KFG01321.1 MEME-1

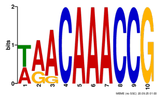
KFG01321.1 MEME-2

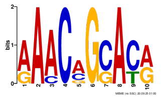
KFG01321.1 MEME-3

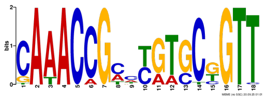
KPC71260.1 MEME-1

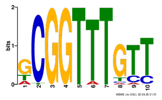
KPC71260.1 MEME-2

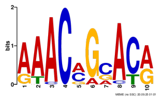
KPC71260.1 MEME-3

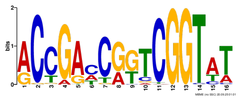
OAL13296.1 MEME-1

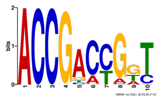
OAL13296.1 MEME-2

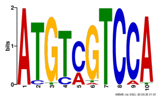
OAL13296.1 MEME-3

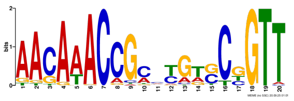
OEJ22426.1 MEME-1

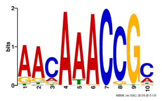
OEJ22426.1 MEME-2

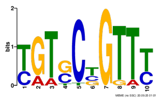
OEJ22426.1 MEME-3

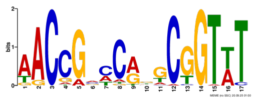
AKJ15784.1 MEME-1

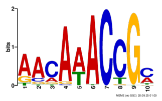
AKJ15784.1 MEME-2

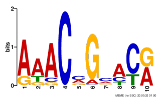
AKJ15784.1 MEME-3

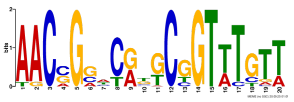
NEC74582.1 MEME-1

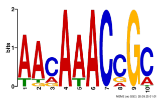
NEC74582.1 MEME-2

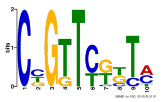
NEC74582.1 MEME-3

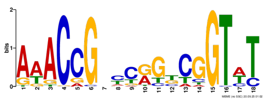
SCK31614.1 MEME-1

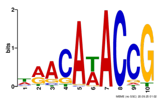
SCK31614.1 MEME-2

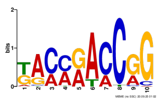
SCK31614.1 MEME-3

#### Group 3

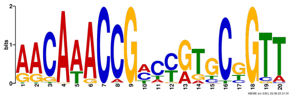

KPC82161.1 MEME-1

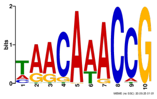

KPC82161.1 MEME-2

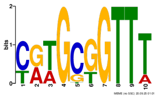

KPC82161.1 MEME-3

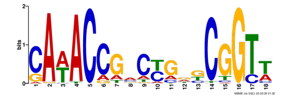

SFL17788.1 MEME-1

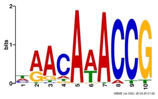

SFL17788.1 MEME-2

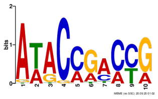

SFL17788.1 MEME-3

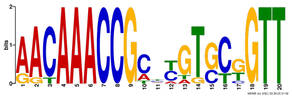

SNX76790.1 MEME-1

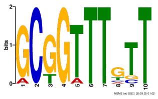

SNX76790.1 MEME-2

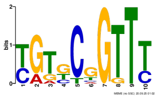

SNX76790.1 MEME-3

#### Group 5

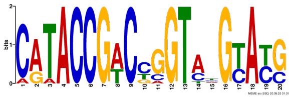

KOU60811.1 MEME-1

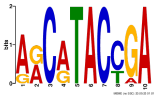

KOU60811.1 MEME-2

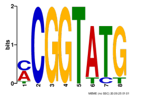

KOU60811.1 MEME-3

#### Group 6

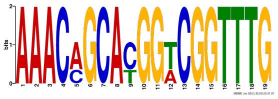

KOV97487.1 MEME-1

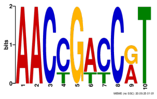

KOV97487.1 MEME-2

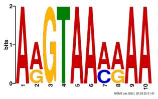

KOV97487.1 MEME-3

#### Group 11

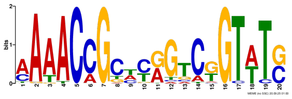

AAX97699.1 MEME-1

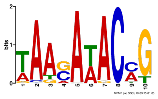

AAX97699.1 MEME-2

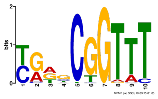

AAX97699.1 MEME-3

#### Group 12

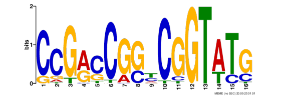
MYU20815.1 MEME-1

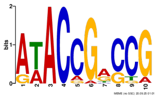
MYU20815.1 MEME-2

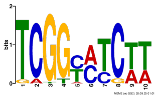
MYU20815.1 MEME-3

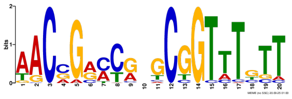
ARI53501.1 MEME-1

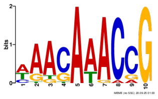
ARI53501.1 MEME-2

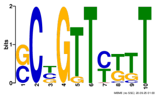
ARI53501.1 MEME-3

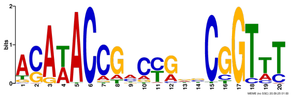
BAA08617.1 MEME-1

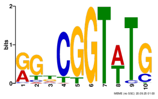
BAA08617.1 MEME-2

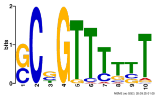
BAA08617.1 MEME-3

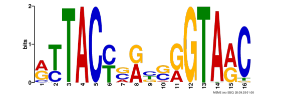
EWS92951.1 MEME-1

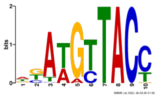
EWS92951.1 MEME-2

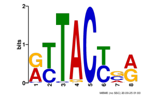
EWS92951.1 MEME-3

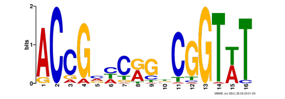
GFN05338.1 MEME-1

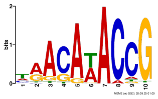
GFN05338.1 MEME-2

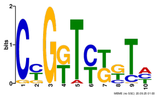
GFN05338.1 MEME-3

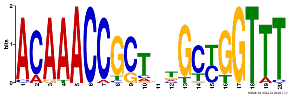
OCC11779.1 MEME-1

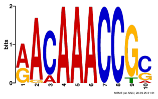
OCC11779.1 MEME-2

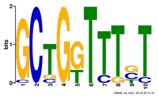
OCC11779.1 MEME-3

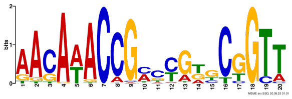
POG46644.1 MEME-1

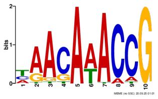
POG46644.1 MEME-2

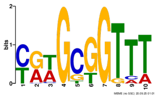
POG46644.1 MEME-3

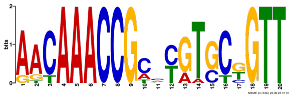
QKW44091.1 MEME-1

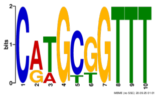
QKW44091.1 MEME-2

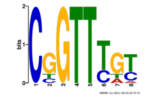
QKW44091.1 MEME-3

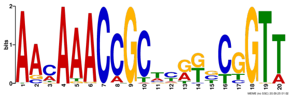
ROV66739.1 MEME-1

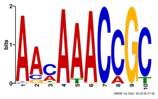
ROV66739.1 MEME-2

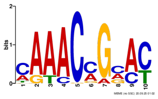
ROV66739.1 MEME-3

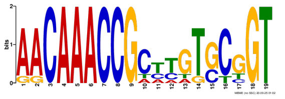
SCD40303.1 MEME-1

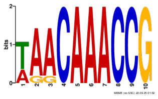
SCD40303.1 MEME-2

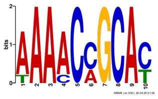
SCD40303.1 MEME-3

#### Group 13

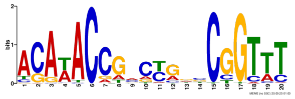

AZS74631.1 MEME-1

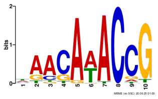

AZS74631.1 MEME-2

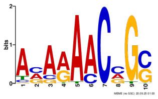

AZS74631.1 MEME-3

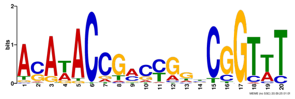

KOG38540.1 MEME-1

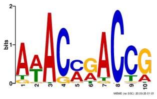

KOG38540.1 MEME-2

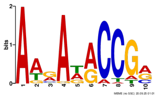

KOG38540.1 MEME-3

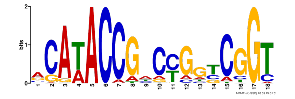

KOT60059.1 MEME-1

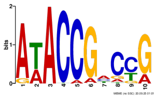

KOT60059.1 MEME-2

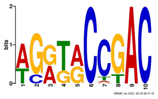

KOT60059.1 MEME-3

### Group 15

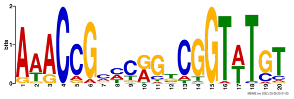
AGK75256.1 MEME-1

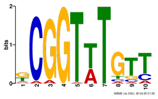
AGK75256.1 MEME-2

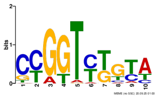
AGK75256.1 MEME-3

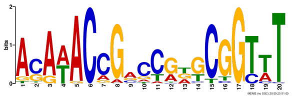
CAA07628.1 MEME-1

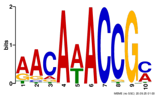
CAA07628.1 MEME-2

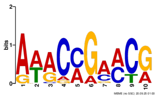
CAA07628.1 MEME-3

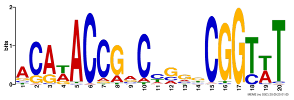
GFN09598.1 MEME-1

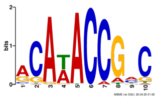
GFN09598.1 MEME-2

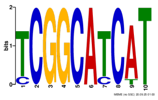
GFN09598.1 MEME-3

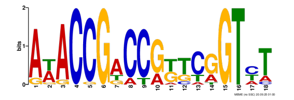
KFK85162.1 MEME-1

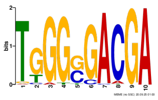
KFK85162.1 MEME-2

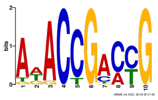
KFK85162.1 MEME-3

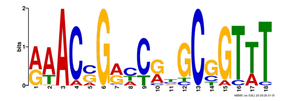
PWS40737.1 MEME-1

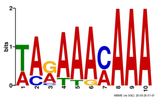
PWS40737.1 MEME-2

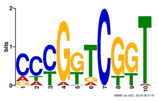
PWS40737.1 MEME-3

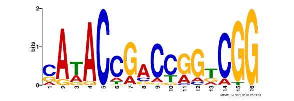
QKW41182.1 MEME-1

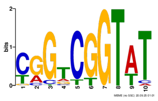
QKW41182.1 MEME-2

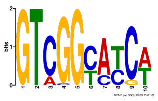
QKW41182.1 MEME-3

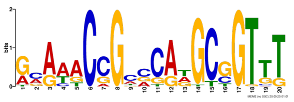
RAN21296.1 MEME-1

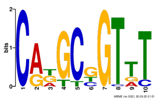
RAN21296.1 MEME-2

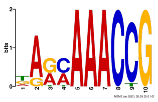
RAN21296.1 MEME-3

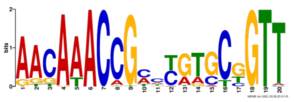
RDL05033.1 MEME-1

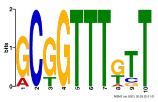
RDL05033.1 MEME-2

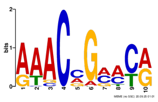
RDL05033.1 MEME-3

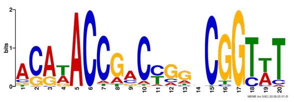
RDV51287.1 MEME-1

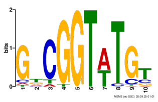
RDV51287.1 MEME-2

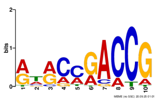
RDV51287.1 MEME-3

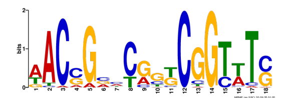
RPK79789.1 MEME-1

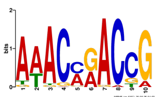
RPK79789.1 MEME-2

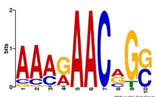
RPK79789.1 MEME-3

#### Group 24

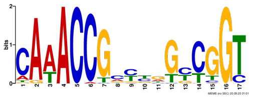

KMS67984.1 MEME-1

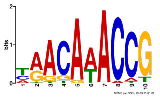

KMS67984.1 MEME-2

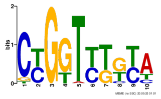

KMS67984.1 MEME-3

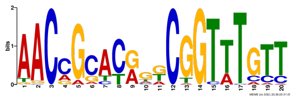

KOU57728.1 MEME-1

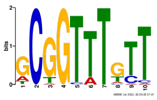

KOU57728.1 MEME-2

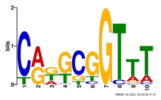

KOU57728.1 MEME-3

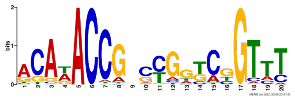

KUN15180.1 MEME-1

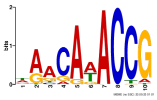

KUN15180.1 MEME-2

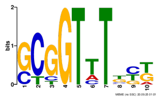

KUN15180.1 MEME-3

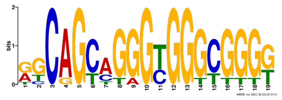

KUN69010.1 MEME-1

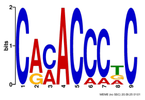

KUN69010.1 MEME-2

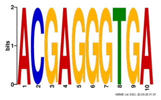

KUN69010.1 MEME-3

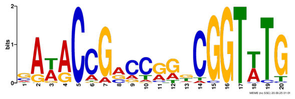

OOQ46925.1 MEME-1

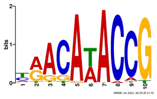

OOQ46925.1 MEME-2

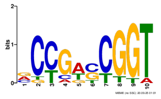

OOQ46925.1 MEME-3

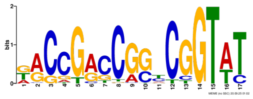

TKT10625.1 MEME-1

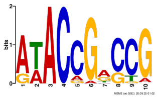

TKT10625.1 MEME-2

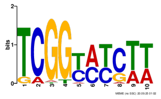

TKT10625.1 MEME-3

### Group 28

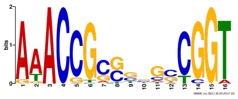
CCA59254.1 MEME-1

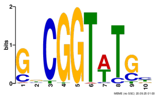
CCA59254.1 MEME-2

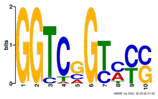
CCA59254.1 MEME-3

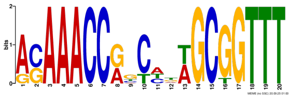
EGX55851.1 MEME-1

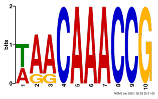
EGX55851.1 MEME-2

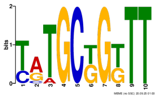
EGX55851.1 MEME-3

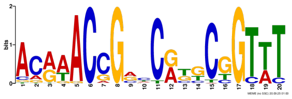
EPJ38720.1 MEME-1

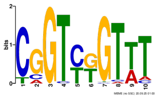
EPJ38720.1 MEME-2

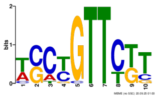
EPJ38720.1 MEME-3

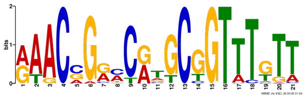
KDN77060.1 MEME-1

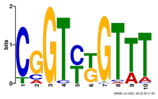
KDN77060.1 MEME-2

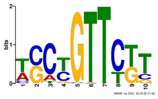
KDN77060.1 MEME-3

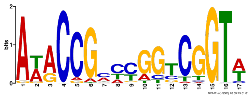
KOT32562.1 MEME-1

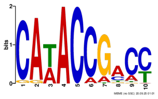
KOT32562.1 MEME-2

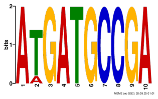
KOT32562.1 MEME-3

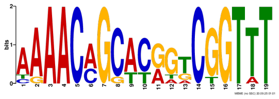
QDI69910.1 MEME-1

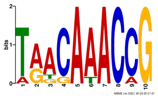
QDI69910.1 MEME-2

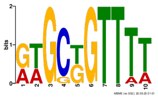
QDI69910.1 MEME-3

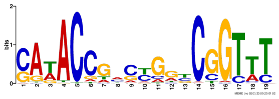
SOR80526.1 MEME-1

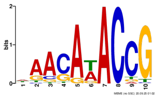
SOR80526.1 MEME-2

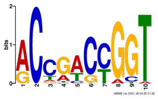
SOR80526.1 MEME-3

### Group 29

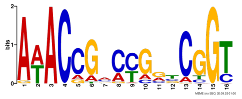
AAK07686.1 MEME-1

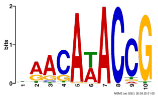
AAK07686.1 MEME-2

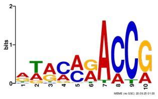
AAK07686.1 MEME-3

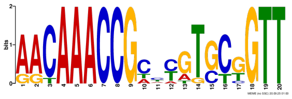
AIS01046.1 MEME-1

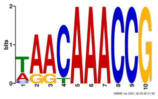
AIS01046.1 MEME-2

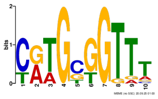
AIS01046.1 MEME-3

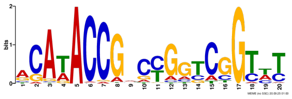
AVZ77953.1 MEME-1

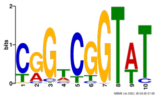
AVZ77953.1 MEME-2

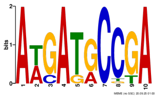
AVZ77953.1 MEME-3

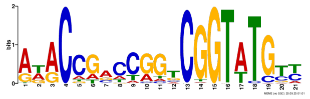
KOU40412.1 MEME-1

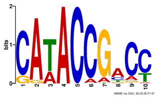
KOU40412.1 MEME-2

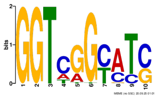
KOU40412.1 MEME-3

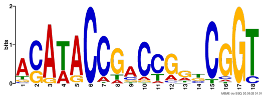
KOY54667.1 MEME-1

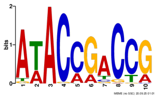
KOY54667.1 MEME-2

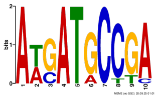
KOY54667.1 MEME-3

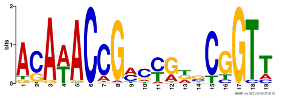
OXY87760.1 MEME-1

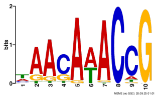
OXY87760.1 MEME-2

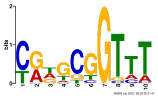
OXY87760.1 MEME-3

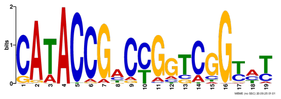
QER87669.1 MEME-1

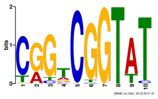
QER87669.1 MEME-2

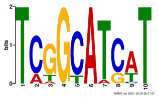
QER87669.1 MEME-3

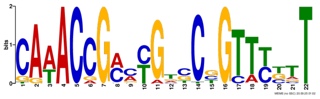
ROP96272.1 MEME-1

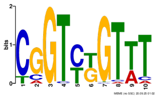
ROP96272.1 MEME-2

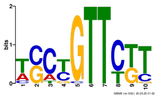
ROP96272.1 MEME-3

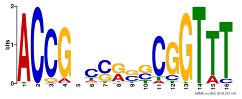
BAC66444.1 MEME-1

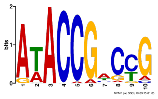
BAC66444.1 MEME-2

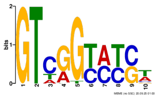
BAC66444.1 MEME-3

#### Group 38

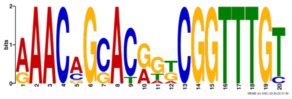
AOR36460.1 MEME-1

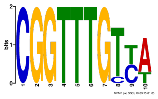
AOR36460.1 MEME-2

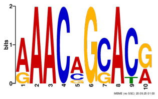
AOR36460.1 MEME-3

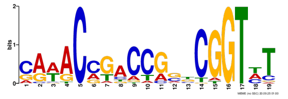
BBC98063.1 MEME-1

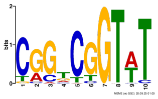
BBC98063.1 MEME-2

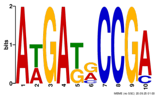
BBC98063.1 MEME-3

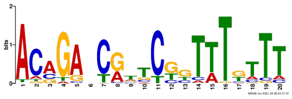
KOG75745.1 MEME-1

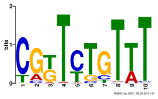
KOG75745.1 MEME-2

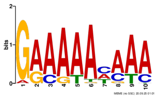
KOG75745.1 MEME-3

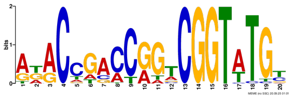
KOV96150.1 MEME-1

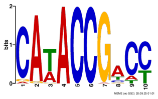
KOV96150.1 MEME-2

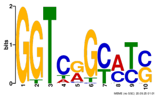
KOV96150.1 MEME-3

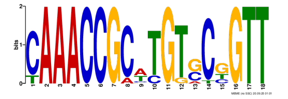
OIJ87288.1 MEME-1

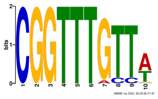
OIJ87288.1 MEME-2

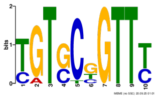
OIJ87288.1 MEME-3

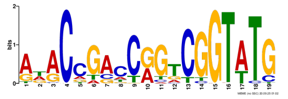
RRQ86746.1 MEME-1

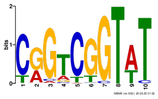
RRQ86746.1 MEME-2

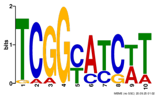
RRQ86746.1 MEME-3

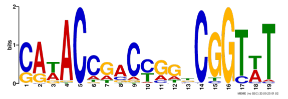
TXJ76008.1 MEME-1

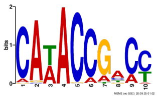
TXJ76008.1 MEME-2

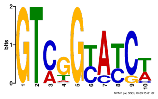
TXJ76008.1 MEME-3

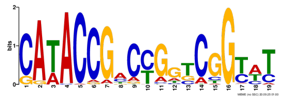
BAC76540.2 MEME-1

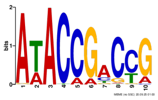
BAC76540.2 MEME-2

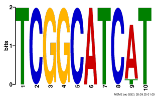
BAC76540.2 MEME-3

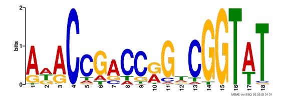
QNT98184.1 MEME-1

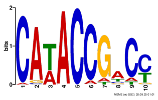
QNT98184.1 MEME-2

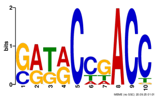
QNT98184.1 MEME-3

#### Group 40

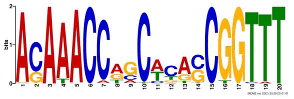
KOG56047.1 MEME-1

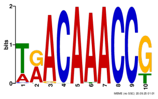
KOG56047.1 MEME-2

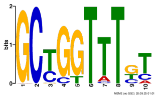
KOG56047.1 MEME-3

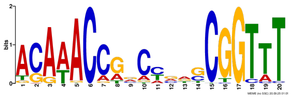
KOU34879.1 MEME-1

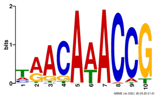
KOU34879.1 MEME-2

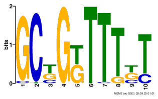
KOU34879.1 MEME-3

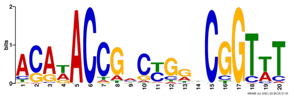
KOU62428.1 MEME-1

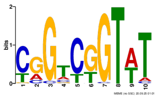
KOU62428.1 MEME-2

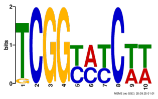
KOU62428.1 MEME-3

#### Group 41

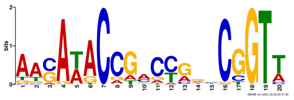
AGN74906.1 MEME-1

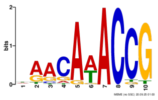
AGN74906.1 MEME-2

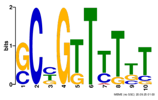
AGN74906.1 MEME-3

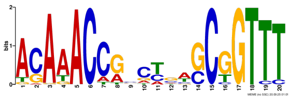
KOV58519.1 MEME-1

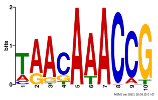
KOV58519.1 MEME-2

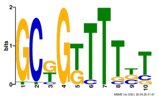
KOV58519.1 MEME-3

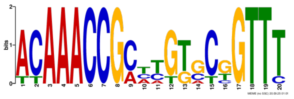
KUN00475.1 MEME-1

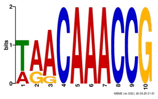
KUN00475.1 MEME-2

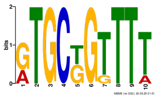
KUN00475.1 MEME-3

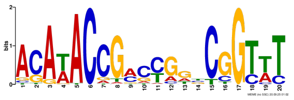
RWZ75665.1 MEME-1

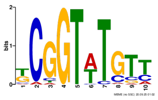
RWZ75665.1 MEME-2

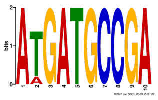
RWZ75665.1 MEME-3

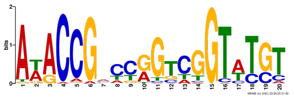
KFG71577.1 MEME-1

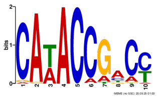
KFG71577.1 MEME-2

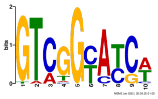
KFG71577.1 MEME-3

#### Group 45

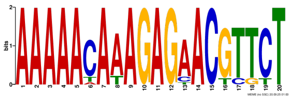
AEW94536.1 MEME-1

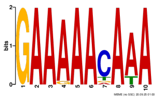
AEW94536.1 MEME-2

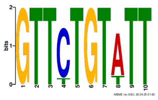
AEW94536.1 MEME-3

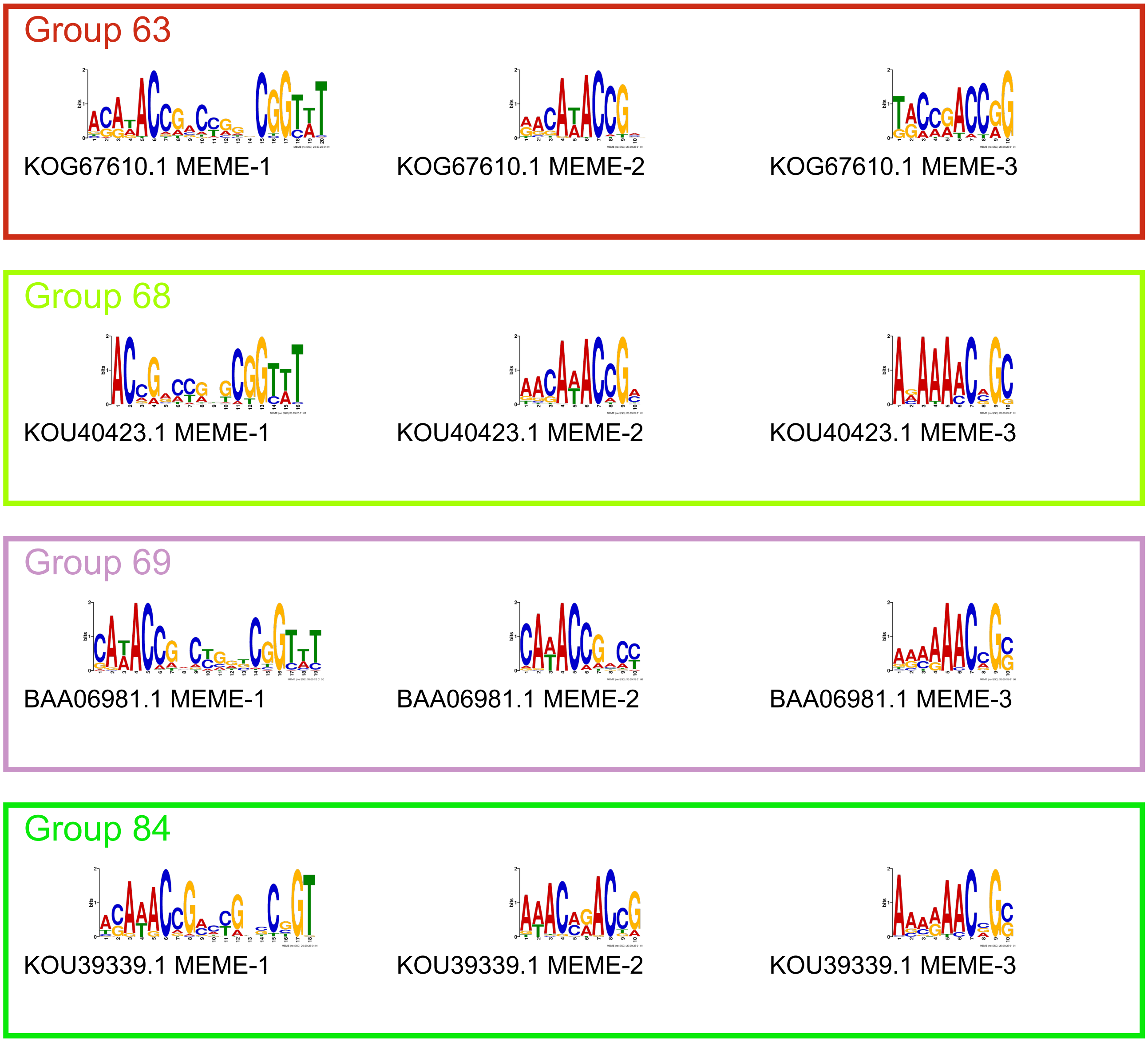

**Supplementary Figure 2.** MEME results for DAP-seq CSR cohort. Group number indicates SSN group of CSR, GenPept ID for the tested CSR is listed alongside the MEME motif number.

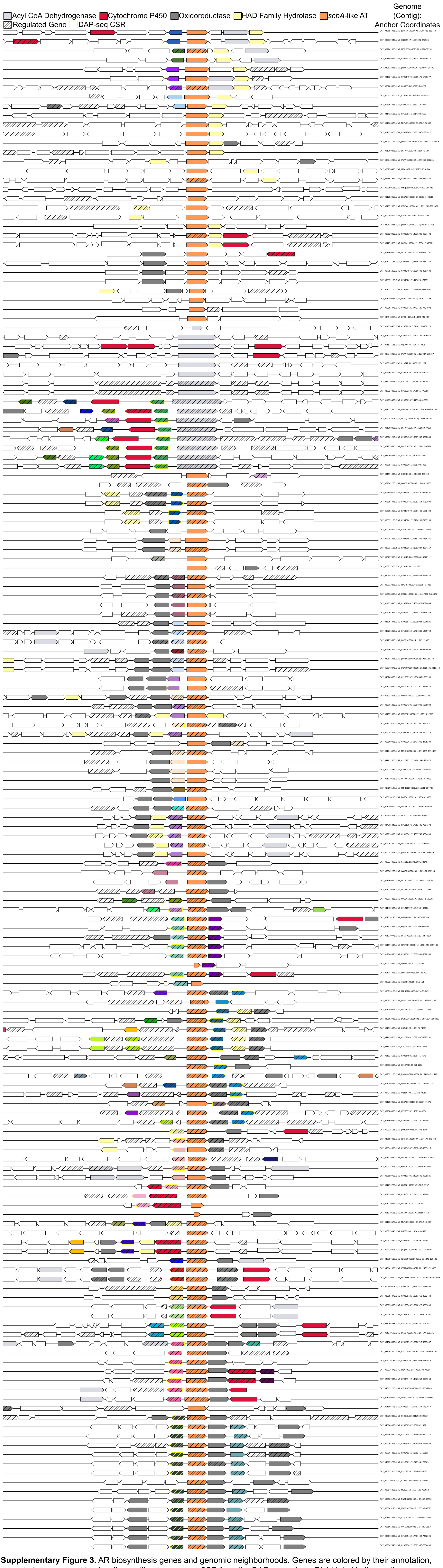

**Supplementary Figure 3.** AR biosynthesis genes and genomic neighborhoods. Genes are colored by their annotation, regulated genes are striped, yellow outlined genes are CSR from the DAP-seq cohort. Right label indicates the genome assembly, contig, and coordinates of anchor.
